# Coherent Gene Assemblies: Example, Yeast Cell Division Cycle, CDC

**DOI:** 10.1101/2021.09.05.459023

**Authors:** Lawrence Sirovich

**Affiliations:** Rockefeller University

## Abstract

A fresh approach to the dynamics of gene assemblies is presented. Central to the exposition are the concepts of: high value genes; correlated activity; and the orderly unfolding of gene dynamics; and especially dynamic mode decomposition, DMD, a remarkable new tool for dissecting dynamics. This program is carried out, in detail, for the Orlando et al yeast database (Orlando et al. 2008).

It is shown that the yeast cell division cycle, CDC, requires no more than a six dimensional space, formed by three complex temporal modal pairs, each associated with characteristic aspects of the cell cycle: (1) A mother cell cohort that follows a fast clock; (2) A daughter cell cohort that follows a slower clock; (3) inherent gene expression, unrelated to the CDC.

A derived set of sixty high-value genes serves as a model for the correlated unfolding of gene activity. Confirmation of our results comes from an independent database, and other considerations. The present analysis, leads naturally, to a Fourier description, for the sparsely sampled data. From this, resolved peak times of gene expression are obtained. This in turn leads to prediction of precise times of expression in the unfolding of the CDC genes. The activation of each gene appears as uncoupled dynamics from the mother and daughter cohorts, of different durations. These deliberations lead to detailed estimates of the fraction of mother and daughter cells, specific estimates of their maturation periods, and specific estimates of the number of genes in these cells.

An algorithmic framework for yeast modeling is proposed, and based on the new analyses, a range of theoretical ideas and new experiments are suggested.

A Supplement contains additional material and other perspectives.

## 1. Introduction

The blueprint of a life form is contained in its DNA genome, as formed from the four bases, [A, C, G, T], as might be assembled in the double helix (Watson and Crick 1953). The genome contains instructions for decoding itself, constructing itself, duplicating itself, and inserting these instructions in the duplicate. This *instruction set* conforms to the Von Neumann (1951) vision of an automaton, an *imitation of life*, also see (Schrödinger 1992). The genetic code (Nirenberg and Matthaei 1961) codes base triplet codons for amino acids, the molecular building blocks of polypeptides. Embedded genomic genes, under *transcription*, synthesize mRNA from DNA, followed by *translation*, the synthesis of protein from mRNA; regarded as the *central dogma* of molecular biology (Crick 1970).

Budding yeast, (Saccharomyces) S. Cerevisiae, a single cell eukaryote, has been well studied using gene arrays (Cho et al. 1998; Orlando et al. 2008; Spellman et al. 1998). Recently, the use of some novel mathematical methods has been applied in the analysis of the yeast CDC (Sirovich 2020), henceforth cited as (I). The present paper significantly refines and extends the previous results. In particular the “co-regulated” genes of (Spellman et al. 1998) are considered, as is a disassembly of the CDC.

In a larger framework, the present analyses might be extended to gene populations that exhibit coherence activity, sometimes dubbed as “pathways or networks “. To avoid misunderstanding, the present goal is the dynamic unfolding, in time of a network or pathway, in contrast to the dynamics of individual genes, as for example in (Tyson 1991).

## 2. Yeast

The fate of a budding yeast cell is to divide asymmetrically into a mother and daughter cell. The period of cell division is roughly an hour or two. In their pioneering paper (Spellman et al. 1998) describe several different means by which to assemble a population of quasi-identical daughter cells, for purposes of tracking the dynamics of the CDC. As in (I) we consider the (Orlando et al. 2008) database that uses elutriation to assemble a population of mother and daughter cells. In the cited reference 5716 genes are monitored, by data sampling, at 16 minute intervals, 15 times, covering roughly two cycles of the CDC. This database contains 2 wild type sets, WT1 and WT2 (also referred to here as G1 and G2, and two mutant sets). Each experimental dataset is thus represented by a matrix of 5716 rows and 15 columns, that describes the temporal expression of all genes, denoted by,

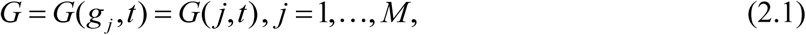

where M = 5716, that specifies the expression levels of each gene, *g*_*j*_, uniformly sampled,

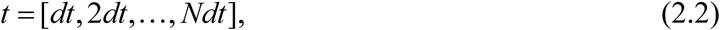

where *N*=15 ×*dt*=16 min.=240 min. Presently, there is no generally accepted set of CDC genes, nor is there an accepted framework for describing the mechanics of the CDC. In (Spellman et al. 1998) the concept of co-regulated genes is proposed as a framework by which to explore related genes of the CDC. One goal of the present study is to shed light on these concepts, solely based on data.

For certain purposes, instead of (2.1) the mean subtracted form,

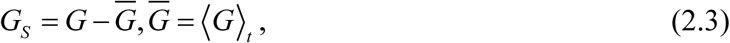

is an effective beginning. *G*_*S*_ is also composed of 15 time snapshots of the 5716 genes. (Since 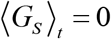, 14 is the correct figure.) The *method of snapshots* (Sirovich 1987) demonstrates that an exact treatment of G_S_ requires no more than fifteen dimensions, as demonstrated by the SVD (Lax 2007), of *G*_*S*_, (2.3),

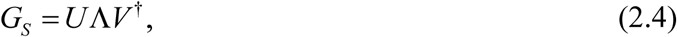

where *U* = _5716_*U*^15^;*V* = _5716_*V*^15^, and Λ is the 15 × 15 diagonal matrix of singular values, and (2.4) is exact. (The 15 × 15 matrix 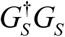 generates Λ and V, from which *U* follows by elementary considerations. The plot below is a log-log plot of the spectrum of variances (squares of singular values).

One can reasonably conclude from Figure 1, that the analysis can be reduced to the six modes, ahead of the *knee*, or intersection. Henceforth, we consider the induced 6-dimensional representation,

**Figure 1.**
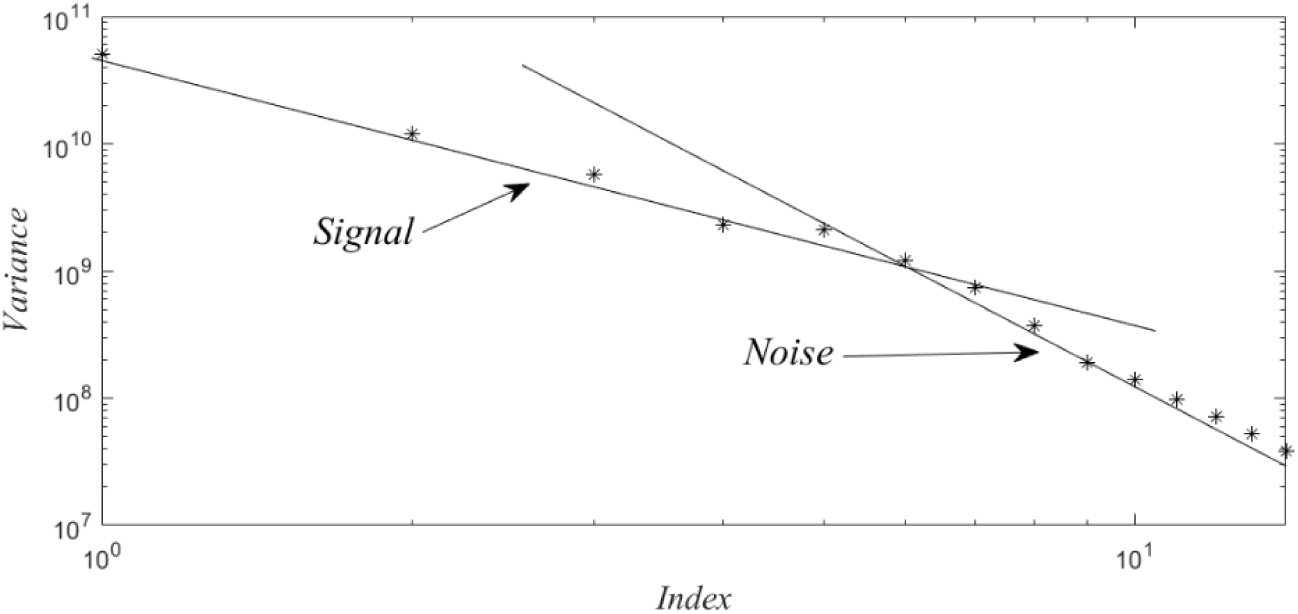
Log-log plots of the variances reveal that these follow two different power laws, which by routine arguments can be associated with signal or noise.

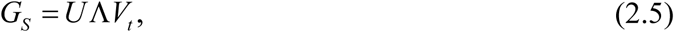

where *V*_*t*_ refer to the 6 × 15 time courses that result from SVD as plotted in Figure 2(A) below.

**Figure 2.**
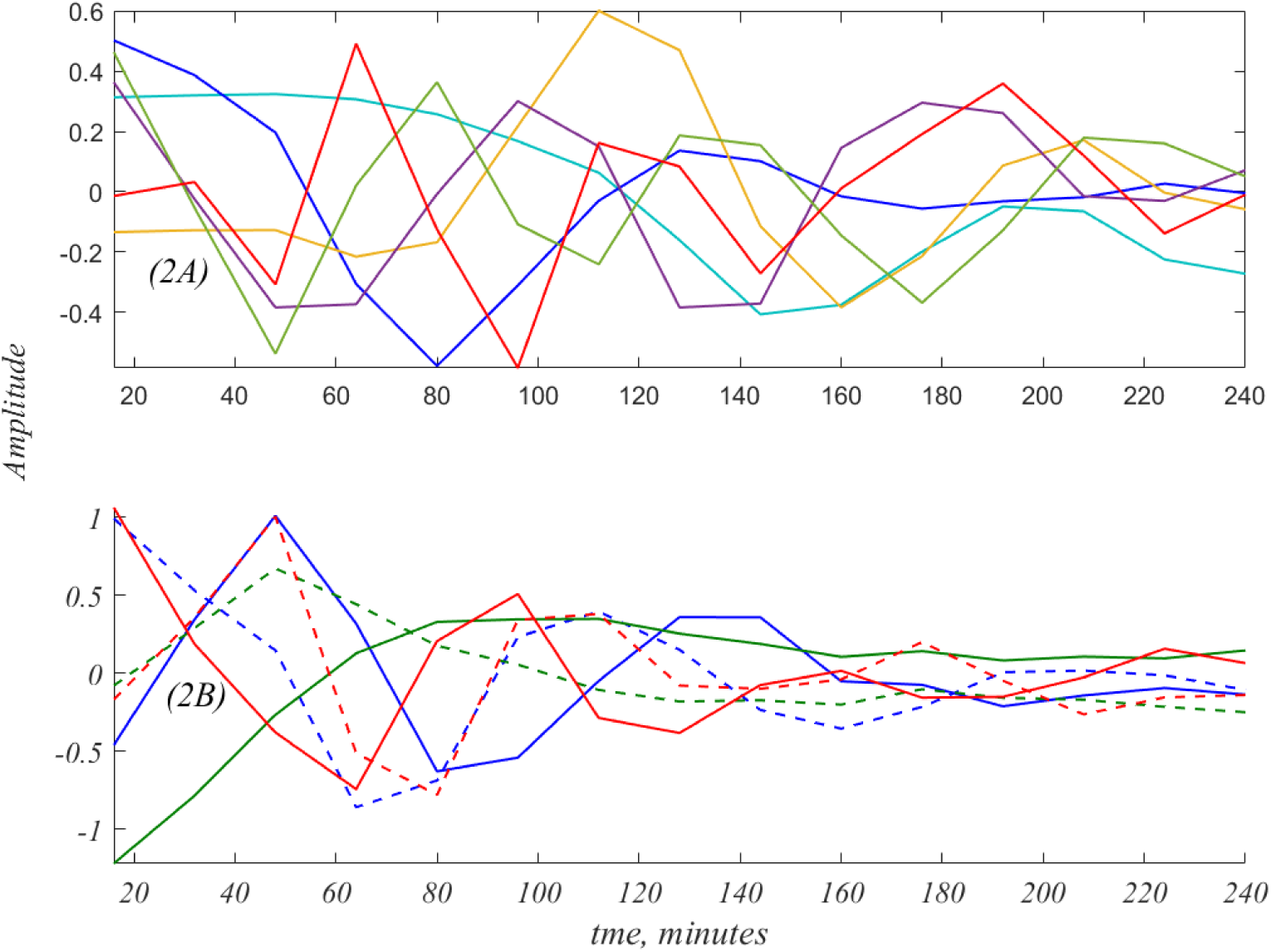
The six time histories of *V*_*t*_ are shown in 2(A). The true underlying dynamics of the data is shown in 2(B), as obtained by Dynamic Mode Decomposition, DMD (Schmid 2010),also see (Kutz et al. 2016) and section 8. This is composed of three complex modes, the real and imaginary parts of which, are depicted by the three colored pairs in 2(B). Further details are given below.

The plots of 2(A) only hint at the actual dynamics. As will be shown these are entangled versions of the intrinsic dynamics, shown in Figure 2(B), see section 8 for the details of the 6×6 matrix, *E*, that disentangles *V*_*t*_, into the three complex modes as shown in Figure 2(B). For the present, note that the decomposition

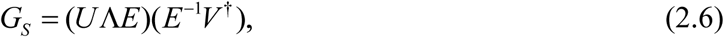

provides a generalization of SVD, (2.4), that is appropriate for data that have a rational ordering of columns, as in dynamics.

### 3. High Value Genes

One can reasonably anticipate two limiting forms of gene expression; steady expression, as might be the case, for the proteins that form the cell wall and membrane; and a briefer activation and later inactivation on a shorter timescale, as might be the case for formation of the cell nucleus and its components. As a criterion for distinguishing these two limits consider the coefficient of variation,

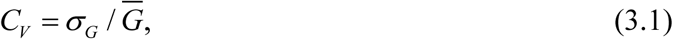

where *σ*_*G*_ denotes the standard deviation of the time history of *G*, and 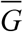, the mean. For example, there are more than 1500 genes for which C_V_<.1 and over 1100 for C_V_>.25. In Table 1 below, the second line specifies six known CDC genes (Orlando et al. 2008), with their coefficient of variation in the line above. Large C_V_ suggests that gene expression is due to transitory mechanisms, as be explained below.

**Table 1.**
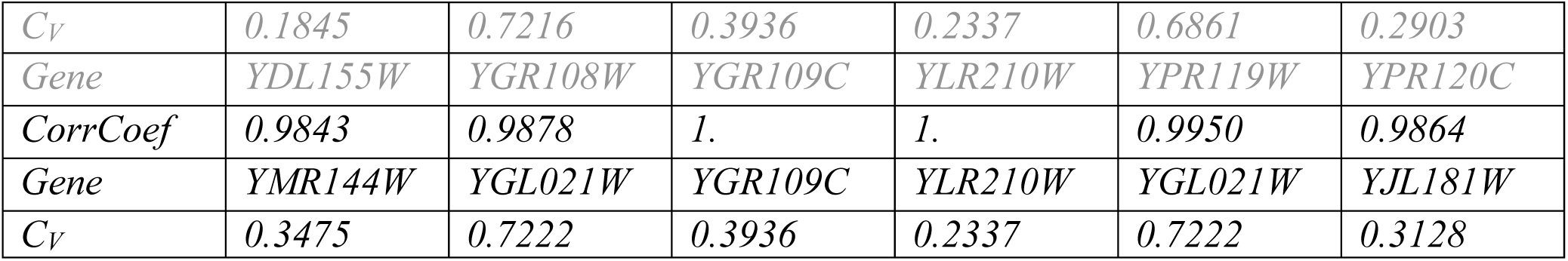
Six known CDC genes (Orlando et al. 2008), line 2. Their coefficients of variation, shown on line 1. Each are highly correlated, line 3, to the genes of line 4; each of which have the higher coefficients of variation, shown on line 5.

Co-regulation implies correlated activity and thee are taken to be linked. The fourth line of the table shows genes highly correlated to those of the second row, shown on the third line, to the genes of the second line. The implication of the Table is that the genes of the fourth line are better gene representatives. Peak times of two like genes are virtually identical.

In general, any gene can be well correlated to many genes. This is illustrated in the next figure the for genes correlated with GR108W, with gene names and coefficients of variation in the legend. *YGL021W* with a C_V_ =*0*.*7222* the best exemplar of this set of highly correlated genes.

As is clear from Figure 3, below, peak locations, ipso facto, must occur at sampling locations, but would be better resolved by interpolation.

**Figure 3.**
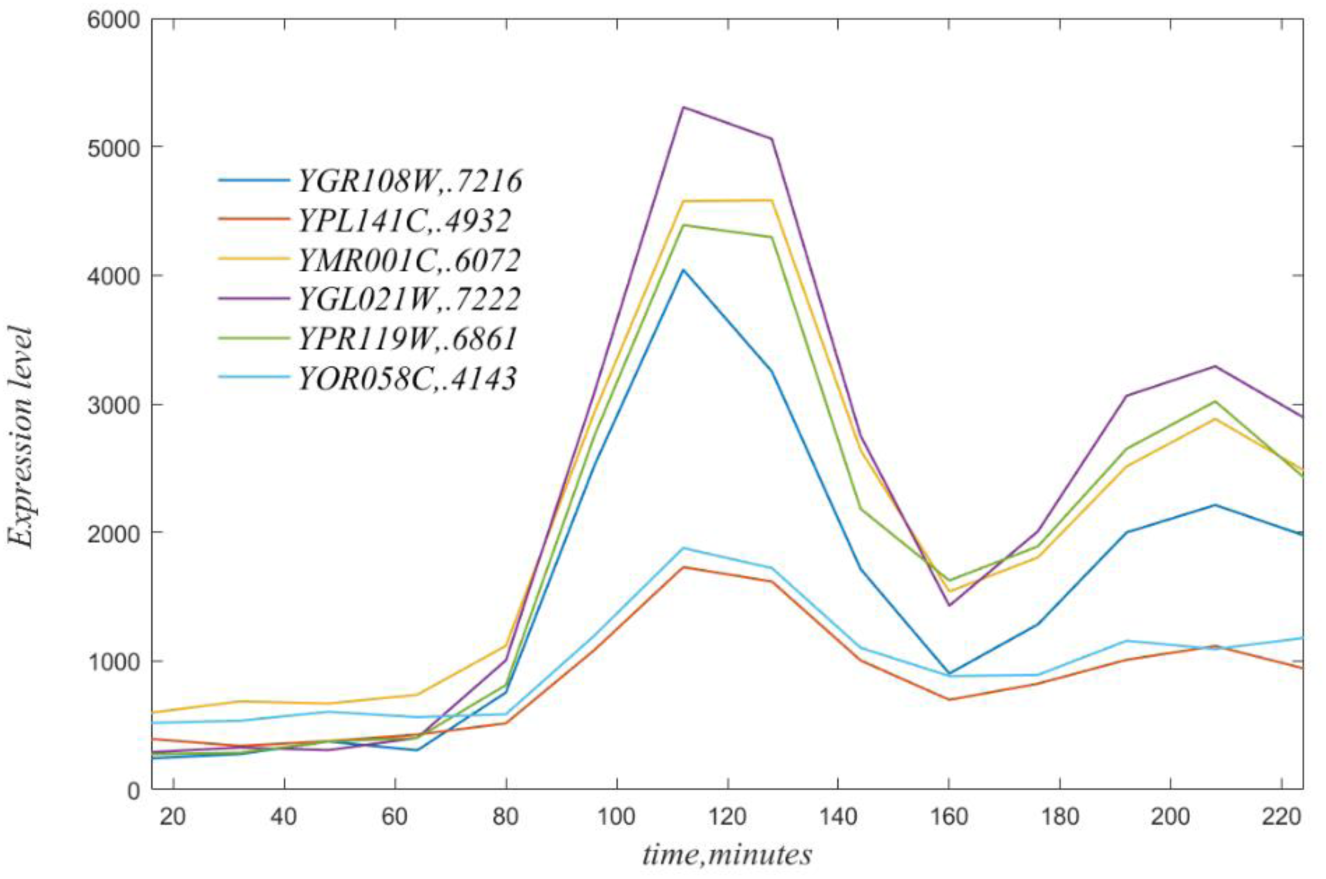
Time courses of genes highly correlated to YGR108W, see legend. Note that each of these has a peak at t = 112, based on the experimental sampling.

#### Gene Selection

As mentioned above there are 1192 genes for which C_V_ >.25. These will be regarded as a starting point for selecting high value genes. A large number exhibit pure exponential decay, starting at extremely high expression values, and are regarded as artefactual. Thus, a second criterion is restriction to time traces that start with relatively small expression, as is the case in Figure 3. For example, a restriction to initial value of a gene at 16 min., of <450, results in a well-correlated set of 109 genes. Figure 4 below shows the correlation image based on the correlation criterion, *ρ* > .85 for the chosen set of 109 genes, in given order.

**Figure 4.**
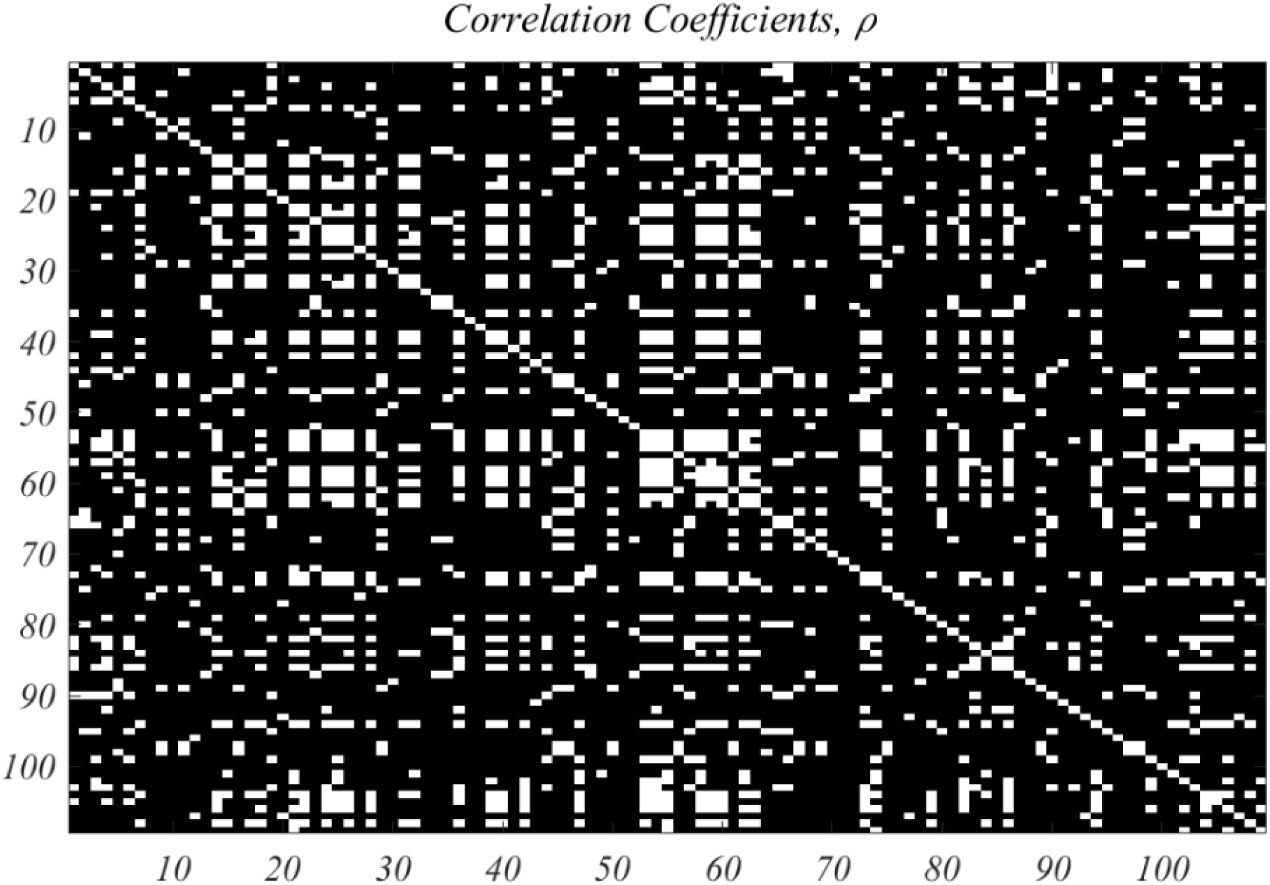
Correlation coefficient of the 109 selected genes, with correlation coefficient, *ρ*, greater than .85 shown as white pixels.

The above figure is based on *r=6* and the 109 high-value genes. The SVD of the derived *G*_*S*_ for this case has the form,

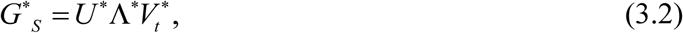

where *U* ^*^ is *109×6*, Λ^*^ = 6× 6 and 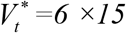. A straightforward calculation shows that this captures more than 99% of the variance of *G*_*S*_. Figure 2 is also based on these choices.

### 4. Analysis of the High Value Genes

The DMD analysis of *G*^*^ produces three complex modes with complex frequencies,

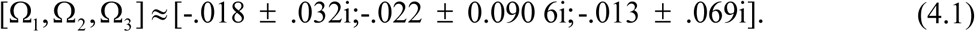

Ω_3_ is identified with daughter cells, with *period T*_3_ = 2*π* / Ω_3_ ≈ 92min.; Ω_2_ is identified with mother cells, with period T_2_ = 2*π* / Ω2 ≈ 71*min* . (smaller daughter cells are well known to take longer to mature.) Depending upon the complexion of the gene sets chosen, there is a range of values for these two periods, however the big picture remains the same. The first period, T_1_, is roughly 200 minutes, close to the duration of the experiment. The time courses following from DMD, and take the form,

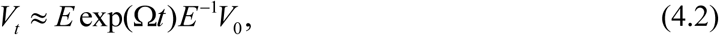

where V_o_ is the initial value, and *E* is derived from the data, and *t* is given by (2.2).The exponential matrix in (4.2) is given by

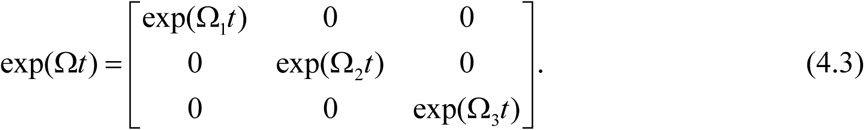

This form speaks volumes. Since each exp(Ω*t*) is a diagonal matrix, these entries are pure exponentials, and thus from (4.2) *E*^−1^*V*_*t*_ disentangles the modes into exponentials. Support for this assertion is shown in Figure 2. On this basis, for data sets of a dynamical type, SVD is generalized by

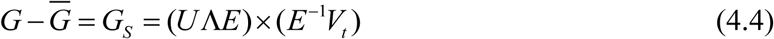

where (*E*^−1^*V*_*t*_), produces un-entangled temporal modes, with amplitudes, (*U*Λ*E*).

And enduring puzzle of SVD analysis has been the origin of the time courses of *V*. This was solved by Peter Schmid (Schmid 2010) with his discovery of DMD, as discussed above. This observation applies, generally, to data of dynamical type, and in particular for our data with the result,

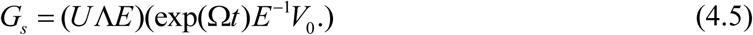

## 4. Unfolding the CDC

A goal here, is the acquisition of a data based model of the CDC. The assumption in this quest is that the CDC is a temporal unfolding of genes, with well-defined activation times. As is clear from Figure 3, time resolution is limited by the coarseness of the sample times. The exponential representation of (4.5) induces a natural Fourier representation that overcomes this limitation. From the calculated frequencies, (4.1), we will *smple* every minute, including the first 16 minutes. In the Figure 3 below, the smooth version of the indicated gene is shown as a dashed curve, and compared with the disentangled experimental modes, (4.4), in order to convey a sense of the Fourier form, as well as the improved peak locations. As an aside, it should be noted that extrapolating backwards in time is rightly regarded as dangerous. However, with a well-defined set of temporal frequencies there is no problem.

Inspection of the 109 highly sampled genes reveals that 43 take on negative values, and 8 have a peak at t = 1. Both conditions are regarded as unacceptable and the corresponding genes will be dropped from consideration. The result is a set of 60 genes that have peak expression times, {*T*_*j*_}, shown in the following figure.

In Figure 7, below, the left image shows the trajectory of gene expression for peak times arranged in ascending order. This compares favorably with the phase plots that appear in (Orlando et al. 2008). At the right is the comparable plot for WT2 under the same gene ordering. It should be noted that the Orlando et al. plots are based on their 440 *consensus* genes. That set contains seven of the present high-value 60 genes, while the co-regulated set of 800 genes in (Spellman et al. 1998), contain 42 of the 60 high-value genes. The table of 60 high-value genes, and their properties appear in the Supplement.

**Figure5.**
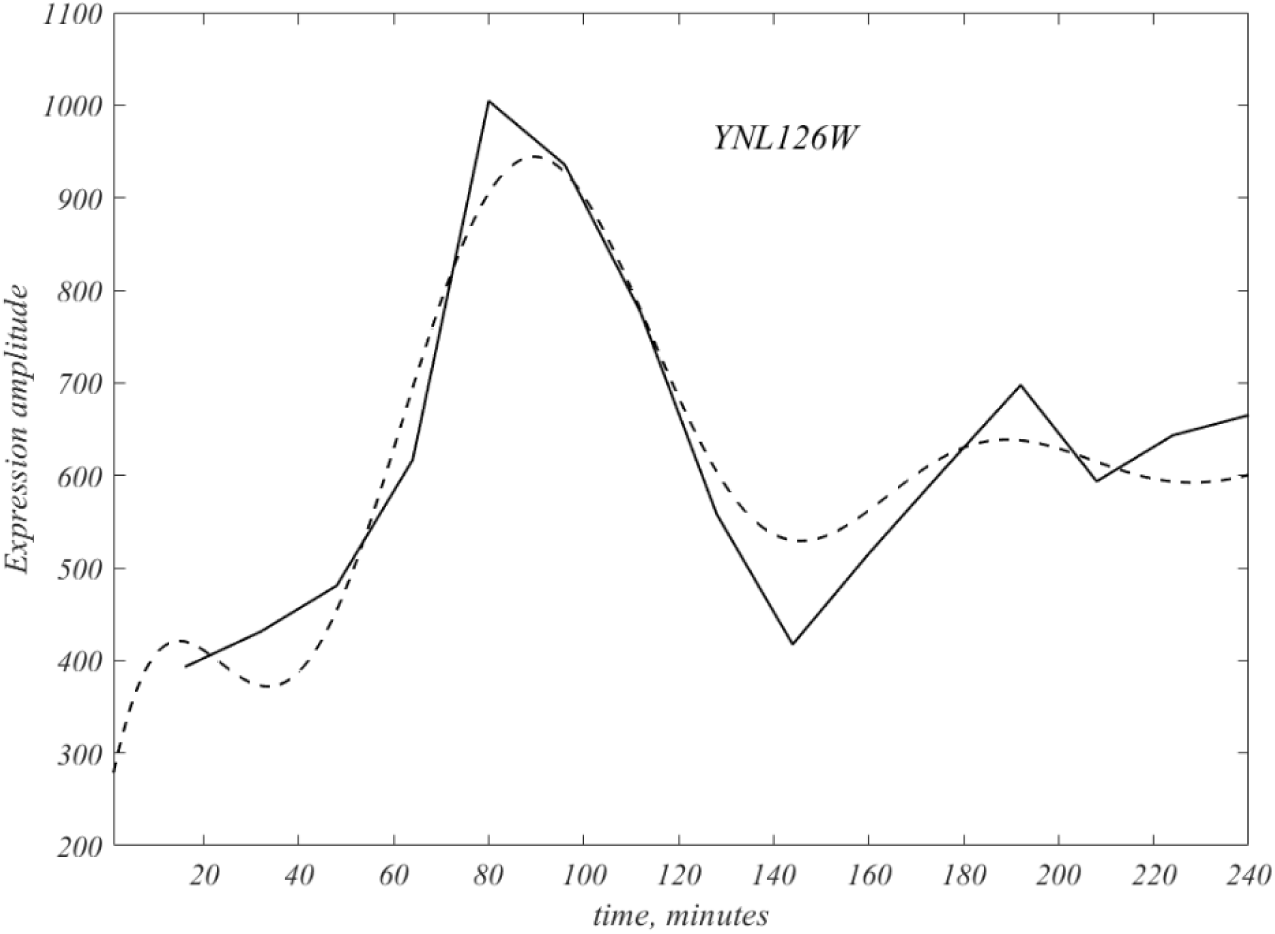
The dashed curve represents the Fourier interpolated version of the original polygonal data of the gene YNL126W, and is the first of the 109 genes. Note the interpolation covers the initial 16 minutes, and also runs over the second attenuated cycle.

**Figure 6.**
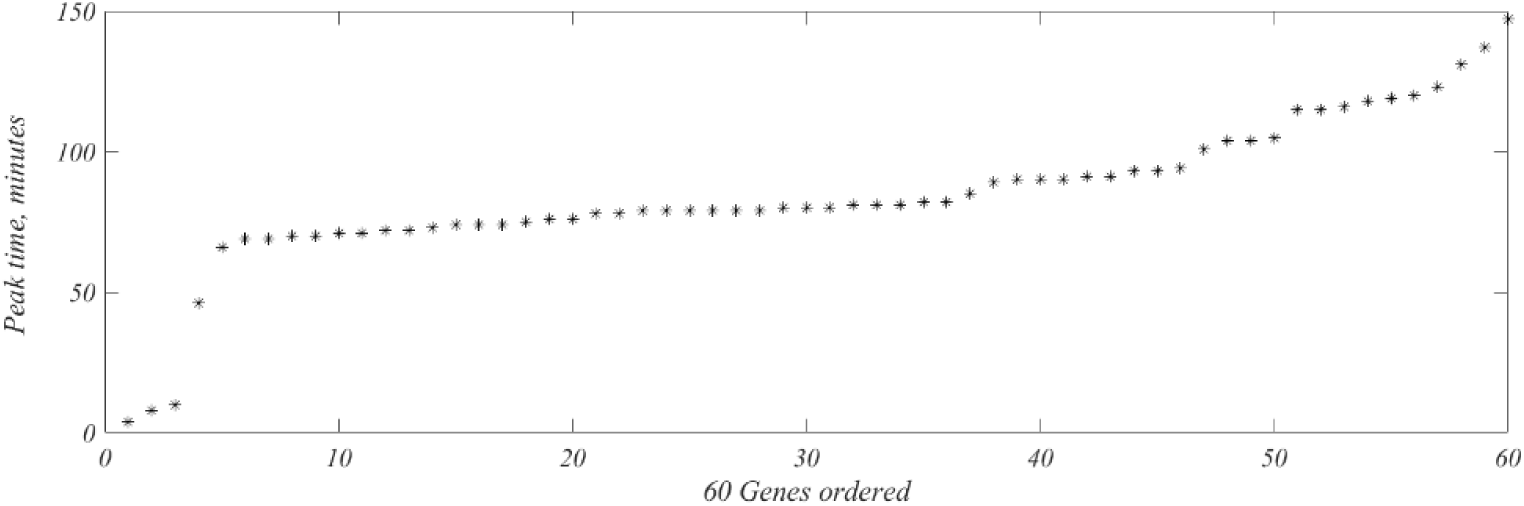
Times, {*T*_*j*_}, of peak appearance, y-axis, of the 60 choice genes.

**Figure 7.**
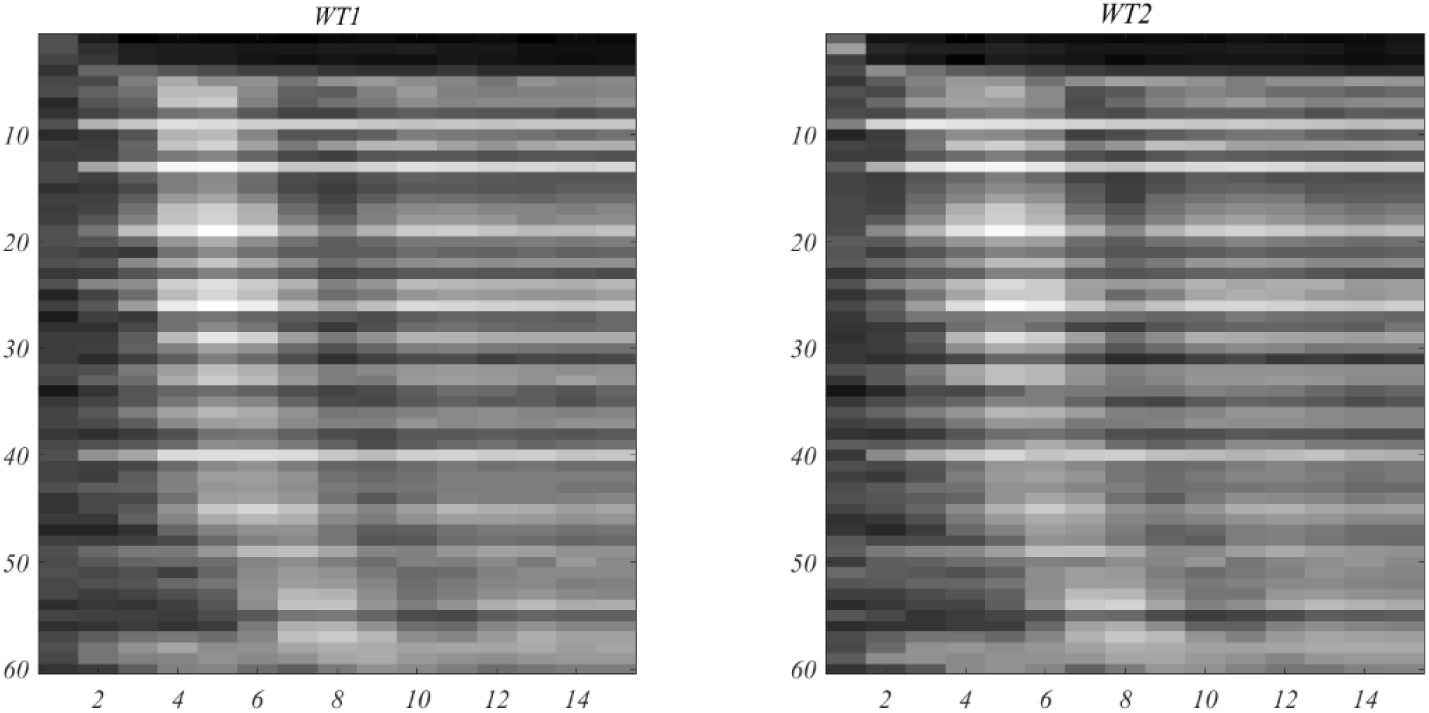
Track of mRNA expression of the 60 selected genes, 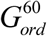, arranged in ascending order of peak activation time. As in Orlando et al., a log transformation has been applied to enhance the image.

### 5. Coherent Gene Sets

Here, *co-regulated* implies *well-correlated*, and we next construct the temporal correlation coefficients. For this purpose, the genes will be ordered as they unfold. This arrangement will be denoted by 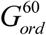. The correlation coefficients of 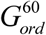, are shown at the left of Figure 8, below.

**Figure 8.**
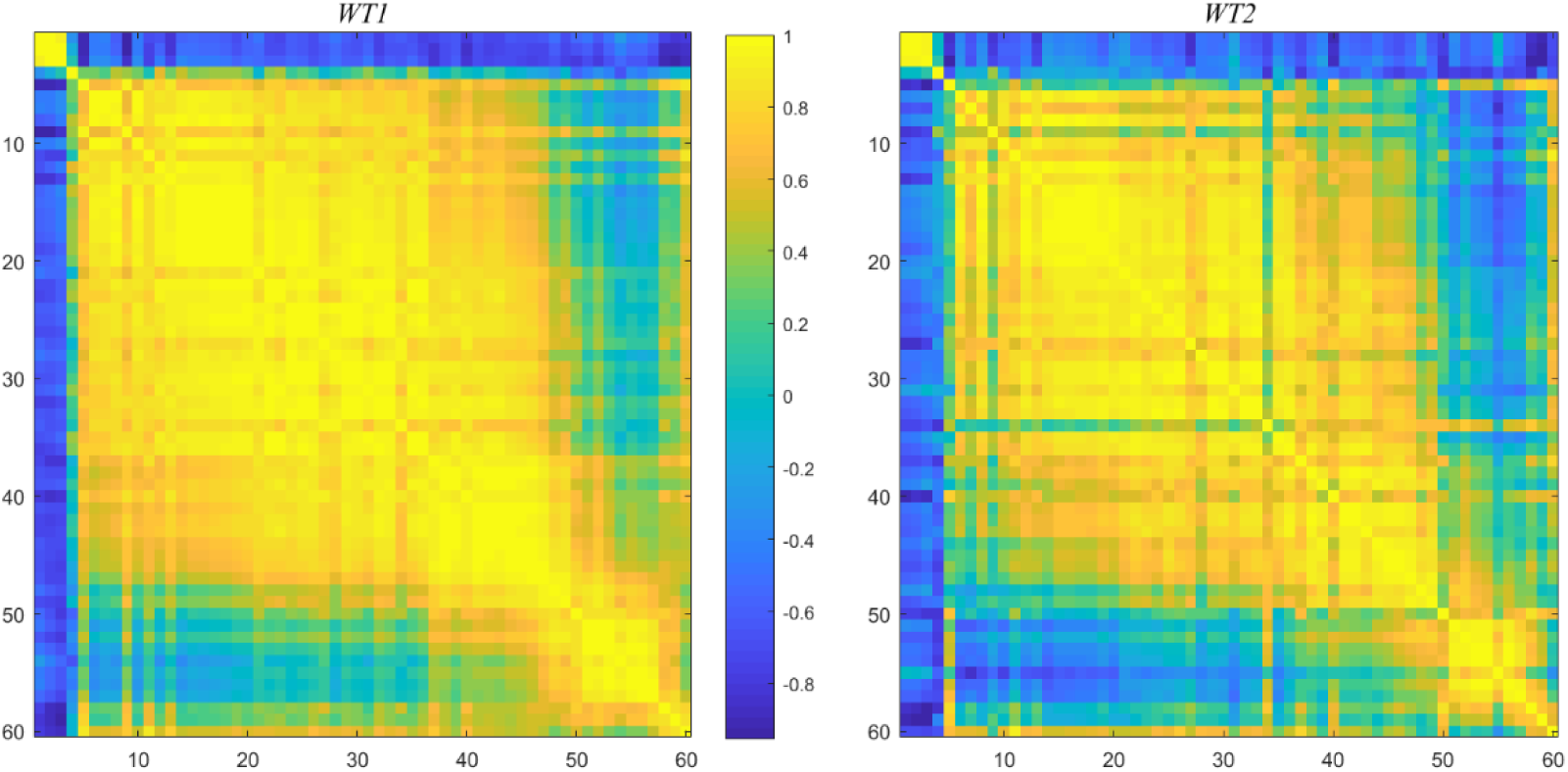
Correlation coefficients, *ρ*, of 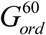, as time ordered, for WT1 and WT2.

The figure on the right is the result of the same calculation, based on the selected ordering applied to WT2. This provides a compelling demonstration that the 60 choice genes are “strongly co-regulated”, in general. A much larger set, 413 genes, similarly constructed, produces an intersection of 204 genes with the Spellman co-regulated set of 800.

Unfolding times is regarded as a reasonable hypothesis for gene ordering; though other possibilities may be considered. For example, ordering the genes in terms of descending correlations, *ρ*, or equivalently increasing temporal distance of time histories. This leads to heat plots and dendrograms, none of which were deemed productive.

### 6. The Single Yeast Cell

Figure 3, exhibits a typical gene time course, and displays single gene expression duration over many tens of minutes. However, the accepted estimate for the duration of gene transcription and translation is 1 to 2 minutes (Milo and Phillips 2015), and for convenience this figure is taken to be one minute in the calculations performed below. To explain what might appear to be an inconsistency, we review the data acquisition procedure, and as will be seen is due to the different maturation periods of mother and daughter cells, and randomness.

In experiments, after assembly of a suitable pool of yeast cells, aliquot removals along with genetic snapshots are obtained at 16 minute intervals, and repeated 15 times. The result is the report of mRNA expression for each gene at each sampling instant. According to (Orlando et al. 2008) each sampling, contains more than 200 yeast cells. To obtain a sense of the process consider gene YGR174 –A, the earliest activated gene, of the 60 gene set, as deconstructed in Figure 9 below.

**Figure 9.**
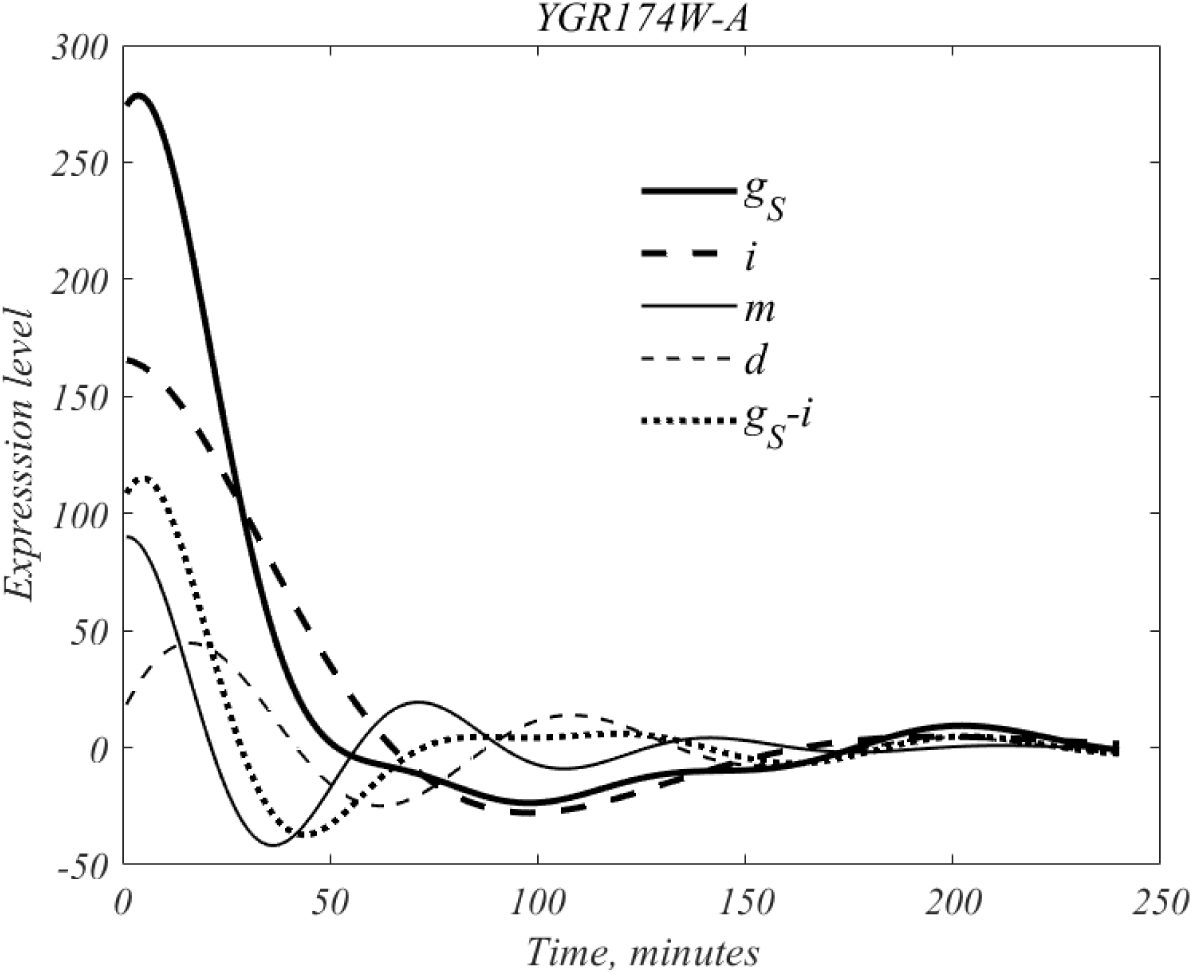
The dynamical composition of gene *YGR174W–A*, the earliest activated gene. See text.

The mean subtracted form of this genes is denoted by g_S_; DMD produces the four traces related by,

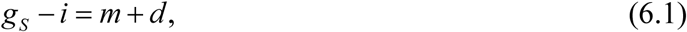

where i, m, d represent the inherent, mother, daughter time courses.

The mother cohort, m, has a peak of 90 at t = 1, of period 71 minutes; and daughter cohort, d, a peak of 45 at 16 minutes and has period 92 minutes. The two entries in the left-hand side of (6.1) are defined over the full interval, and their difference has a peak of 115 at 5 minutes. If we denote by *N*_*M*_ and *N*_*D*_ the unknown number of mother and daughter cells that are participating in the signal, then straightforward back of the envelope calculation, based on peak quantities and locations, shows that *N*_*M*_ / *N*_*D*_ ∼.285, from which the population fraction,

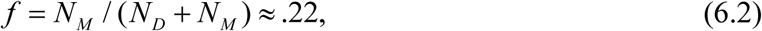

follows. Thus, there are more than three times as many daughter cells as mother cells in an average aliquot.

For formal purposes the inherent signal and the mean will be divided into 2 parts, as follows,

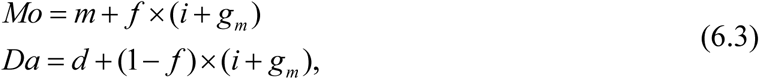

so that the total signal is equals *Mo*+*Da*.

where mother, daughter, inherent and mean our shown in Figure 9. As shown in Figure 10, below, the full signal this gene equals *Mo*+*Da*..

**Figure 10.**
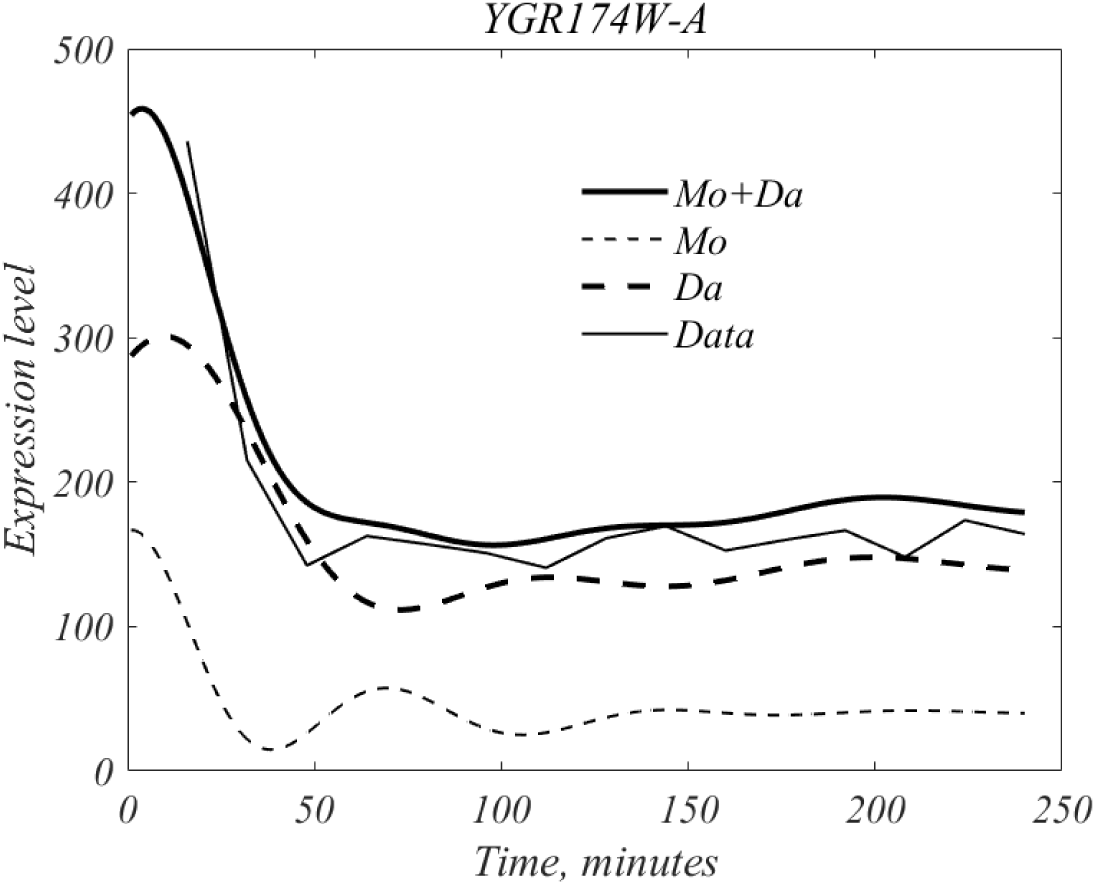
Mother cell dynamics are initiated at roughly t = 0, and daughter cell dynamics at roughly 10 minutes, with each being responsible for a steady production at large t. The two add to yield the overall gene activity shown in the heavy curve.

To summarize, genes are expressed both in a background manner, inherent expression, and in a scheduled manner, to peak at some specific time. It is also clear from this analysis that the activities of mother and daughter cells are not coupled.

#### A Yeast Model

Next, we consider a possible computational model of the yeast cell. For this we focus on the gene traces displayed in Figure 10. While additional genes might be included in the model, the uncertainties in experimental results (Gerstein et al. 2007; Milo and Phillips 2015) do not justify such generalizations. Our purpose in this exercise is merely to demonstrates that a practical framework can be created.

To start, it is noted that estimates of protein molecules per yeast cell are in the range of ∼ 5×10^7^ (Milo and Phillips 2015). Since there are roughly 6000 genes in the yeast cell under consideration, the estimated number of such molecules per gene is ∼ 5×10^7^ / 6000 ≈ 8.3×10^3^. This is a *ballpark* figure, and in this spirit other considerations such as protein degradation will be ignored. Further, the duration of proteins expression will be taken as 1 minute. If we denote the mother and daughter dynamics of Figure 10 by *m*(*t*)&*d*(*t*), respectively, the number of proteins at maturation for mother and daughter cells is estimated from by,

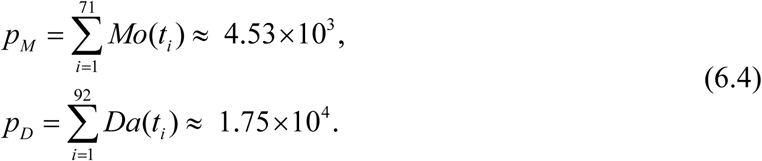

The ratio p_D_/p_M_ =.26 is remarkably close to the above mentioned *N*_*M*_ / *N*_*D*_ ∼.285 . Since these estimates are in the range of the ballpark gene estimate of 8.3×10^3^, it is surmised that the Orlando et al data was normalized by the estimated number of cells in an aliquots, >200. These considerations also suggest that cell size differences of mother and daughter cells plays little or no role in protein content at maturation.

On the basis of these deliberations one might contemplate creating an algorithmic model of the CDC. Randomization can be introduced through variations in mother and daughter CDC periods, and variations in the number of mother and daughter cells, say adding up to roughly 200. This is a future project, which can useful only with better knowledge and precision of the quantities involved.

### 7. Additional Comments

The high degree of correlation, seen in Figure 6 tells little beyond timing. For example, it does not imply anything certain about gene interactions, nor is there any information about the activation and deactivation times of genes. The mechanism by which budding yeast cell assembles itself, is an open question. Since no outside intervention is in play, it is noncontroversial to say, that the cell self-assembles. Just how this self-assembly takes place is another open question, it might e.g. only be a matter of proteins falling into their proper place, based on the timing of gene expression. In this case the cell model is an *assembly line* for proteins that arrive in an orderly fashion.

In an effort to introduce some additional theory it is noted activation and inactivation of gene expression may be likened to an equilibrium disturbance, followed by restoration, (suggestive of wave phenomena.). In this connection it is noted that the mother and daughter modes that describe the CDC each follow oscillator dynamics (Sirovich 2020)

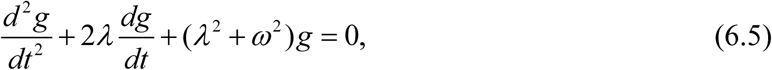

which has solutions in the form,

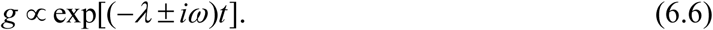

In this connection, we can define a new variable, *v*, by,

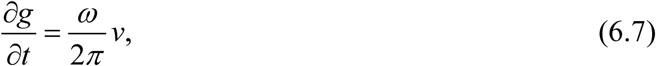

which since 2*π* / *ω* gives the period of mRNA, might be reasonably associated with rate of protein production, and from (6.5) *v* is governed by

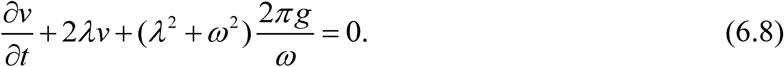

T3he pair of equations (6.7) & (6.8) may be viewed as a coarse-grained version of the *central dogma*.

As mentioned the dynamics of transcription and translation occurs on a scale of ∼ minute, or so. As dictated by the experiments, our deliberations are based on timescales large compared to one minute, and thus, only genes and their proteins can figure in the description. This is consistent with “one gene-one polypeptide” view of (Beadle and Tatum 1941). Impressionistically, this Beadle-Tatum view is a macroscopic description.

A key result of the present investigation is the remarkable ability of DMD, to distinguish dynamic characteristics of the mother and daughter cohorts. Gene experiments typically attempt to sequester daughter cohorts, and it is natural on the basis of the methods pursued here to consider what the outcome would have been of considering complementary populations, and furthermore to monitor the base population, without any form of sequestering. Given the remarkable ability of DMD to parse dynamic activity more complicated population yeast populations might be considered. Hopefully, this ability to distinguish yeast subpopulations will lead to new ways to probe into the yeast life form. Another future goal, is that the algorithmic model touched on here can be further advanced, since a falsifiable model is always desirable.

### 8. Methods

#### Dynamic Mode Decomposition

Traditional signal analysis is based on the hypothesis that a *signal* is an admixture of sinusoids, and therefore that Fourier analysis can decompose the signal into his Fourier components. DMD may be regarded as an extension of this idea. Suppose for real λ & ω, we define a complex frequency,

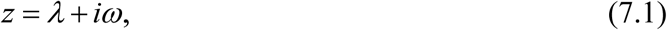

with corresponding complex signal,

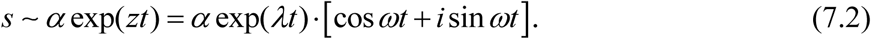

*DMD* can be regarded as a method for extracting such a signal, and more generally, admixtures of such signals. In this respect Fourier analysis is a subset of *DMD*.

Typically, a laboratory signal is a uniformly sampled version of the continuous case. Suppose for example the uniformly sampled times are,

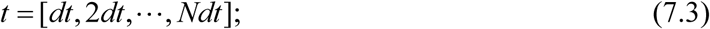

in which case (7.2) becomes the geometric sequence.

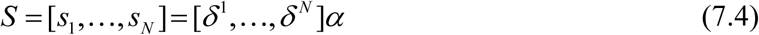

where

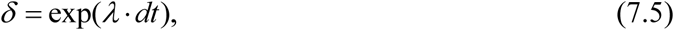

which is therefore the generator of the sampled signal. Reciprocally,, if {*s*_*j*_} is (noisy) data, we can seek a generator, *A*, such that

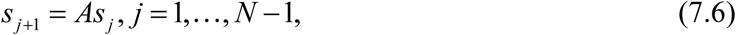

which can be viewed as a case of many equations for the one unknown, *A*. Least squares is the suggested approach in this case. The error in applying (7.6) is

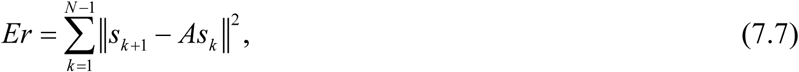

and the (least-squares) minimization of (7.7) produces the solution

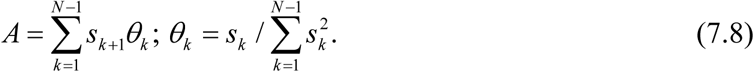

A more compact form is obtained by defining the vectors,

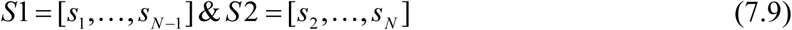

in which terms (7.8) can be written as

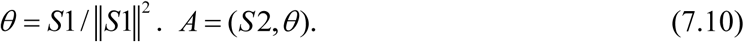

for later purposes observe that if the problem is posed as,

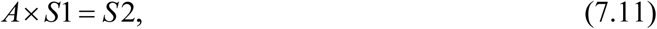

then it is solved by

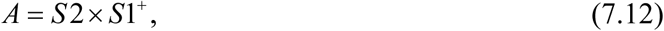

Where *S*1^+^ is referred to as the Moore-Penrose inverse (Golub and Van Loan 1989). Clearly, *S*1^+^ = *θ*, and (7.12) is equivalent to minimizing (7.7). Briefly stated, the Moore Penrose inverse has the smallest norm of the many solutions that solve (7.11).

In the general case, of multiple complex signals, we are confronted by a matrix *V(t)*,

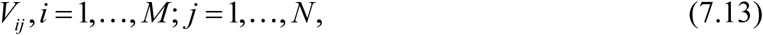

with M time histories, each sampled uniformly N times. For example 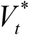 (3.2) is 6 × 15. Under the hypothesis that the data is composed of complex exponentials, one can seek the generator matrix, *A* by defining

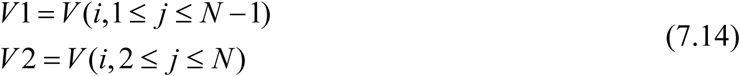

so that

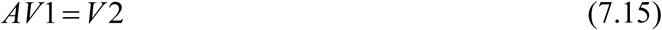

which is solved by the Moore-Penrose inverse, *V*1^+^

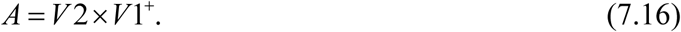

The spectral decomposition of the generator matrix, *A*,

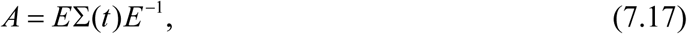

produce complex frequencies as eigenvalues of the diagonal matrix Σ, and that the corresponding eigenvectors in *E*_−_ disentangle the time courses produced by SVD. Evidence supporting both assertions appear in the data analysis. The construction (7.17) permits generalization of SVD, (2.6). If we write

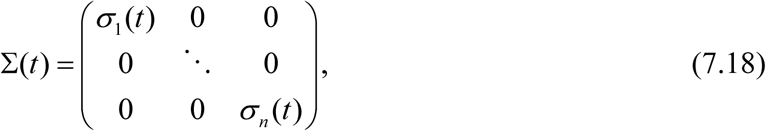

where n = 6 in the yeast application, then it proves to be convenient to transform to the hypothesized exponential form

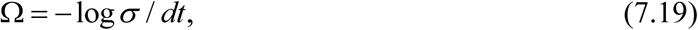

which is just the inverse of (7.5). *dt* is the sampling interval (2.2) and (7.18) transforms to (4.3) In the interest of brevity, we forgo examples. Consideration of synthetic data generated by,

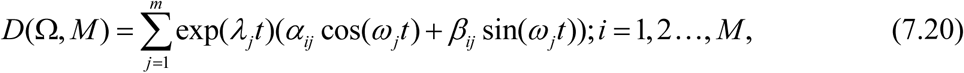

with each of the M trials an admixture of m complex signals and randomly chosen coefficients shows a remarkable accuracy in recovering the frequency content after relatively few trials.

